# Individual vocal signatures show reduced complexity following invasion

**DOI:** 10.1101/2020.09.02.280297

**Authors:** Grace Smith-Vidaurre, Valeria Perez, Timothy F. Wright

## Abstract

The manner in which vocal learning is used for social recognition may be sensitive to the social environment. Biological invaders capable of vocal learning are useful for testing this possibility, as invasion alters population size. If vocal learning is used for individual recognition, then individual identity should be encoded in frequency modulation patterns of acoustic signals. Furthermore, frequency modulation patterns should be more complex in larger social groups, reflecting greater selection for individual distinctiveness. We compared social group sizes and used supervised machine learning and frequency contours to compare contact call structure between native range monk parakeets (*Myiopsitta monachus*) in Uruguay and invasive range populations in the U.S. Invasive range sites exhibited fewer nests and simpler frequency modulation patterns. Beecher’s statistic revealed reduced individual identity content and fewer possible unique individual signatures in invasive range calls. Lower estimated social densities and simpler individual signatures are consistent with relaxed selection on the complexity of calls learned for individual recognition in smaller social groups. These findings run counter to the traditional view that vocal learning is used for imitation, and suggest that vocal learning can be employed to produce individual vocal signatures in a manner sensitive to local population size.

## Introduction

One way in which vocal learning can be used is to signal group identity for social recognition [1–3]. Patterns of acoustic convergence within social groups, consistent with vocal learning being employed for group recognition, have been identified in cetaceans, bats, songbirds, and parrots [3]. Vocal learning may also be used to create individually distinctive acoustic signals, often termed “individual signatures”, as found in many of the same taxonomic groups [1,4–6]. The manner in which different taxa use vocal learning to recognize group members could be sensitive to population size and social dynamics.

In larger populations, greater social density results in more individuals for potential receivers to discriminate, leading to increased uncertainty about signalers’ identities, and increased selection on signalers to produce distinctive individual signatures [7]. In species that employ vocal learning for individual vocal recognition, such selection to produce individually distinctive signals should manifest in the acoustic structure of learned calls. For instance, learning can be employed to produce individually distinctive frequency modulation patterns in acoustic signals [4–6,8]. In such systems, frequency modulation patterns in learned signals should be more complex in larger social groups, and simpler in smaller social groups, leading to fewer potential unique individual signatures [9]. Biological invaders offer useful models for addressing these ideas, as invasive populations often exhibit reduced population sizes compared to the native range [10].

We asked whether the way in which vocal learning is employed for social recognition is resilient or sensitive to changes in population size by evaluating contact calls of an invasive parrot. Monk parakeets (*Myiopsitta monachus*) are native to South America, may use vocal learning for individual recognition [11], and have established invasive populations across the world through the pet trade [12]. We predicted that estimated social densities would be lower following invasion, and frequency modulation patterns in invasive range contact calls would be simpler compared to the native range.

## Methods

### Contact call recording

Native range contact calls were recorded in 2017 at nest sites in Uruguay, as previously described [11]. Invasive range contact calls were recorded at nest sites across five states in the U.S. over different years. When possible, we estimated the numbers of nests visible at recording sites (Supplementary Table 1). We obtained previously published calls recorded in 2004 in Connecticut, Florida, Louisiana, and Texas [13]. Calls were also recorded in Texas and New Orleans in 2011, Arizona in 2018, and Texas in 2019. We repeatedly sampled contact calls from 8 native and 9 invasive birds, otherwise, a single contact call was obtained per unmarked bird (Supplementary Tables 2 - 4). Most recording sessions were performed with Marantz PMD661 MKII and PMD660 solid state recorders, Sennheiser ME67 long shotgun microphones, and digitized at 44100 Hz sampling rate and 16 bit depth. Invasive range 2004 sessions employed Marantz PMD670 or PMD690 recorders with Sennheiser ME67/K6 shotgun microphones, digitized at 48000 Hz and 16 bits [13]. Invasive range calls were selected using Raven version 1.4 [14], and pre-processing was performed with the warbleR package in R to retain high quality calls [15,16].

### Acoustic structure analyses

Differences in call structure between ranges were evaluated with supervised machine learning models that classified calls back to each range. Models were built with 203 predictors, including 15 standard acoustic measurements and 188 features (Supplementary Methods 2.1.1). Spectrum-based measurements were obtained using a Hanning window, window length of 398, window overlap of 90 for Fourier transformations, and a bandpass filter of 0.5 to 9kHz [11]. 1561 calls were split into training, validation, and prediction datasets. 548 calls used for prediction (230 native, 318 invasive) were visualized in two-dimensional acoustic space by applying multidimensional scaling (MDS) to the proximity matrix of the final random forests model. A Gaussian kernel density estimator was applied to MDS coordinates to yield density in random forests acoustic space.

80 calls were subsampled to evaluate frequency modulation patterns between ranges. 10 sites were randomly selected per range, and 4 calls randomly chosen per site. Fundamental frequencies were estimated at 100 timepoints per call, and used to manually trace the second harmonic with warbleR [15]. 5 points were dropped from the start and end of each frequency contour, and spline interpolation performed across the remaining 90 points with smoothing. Smoothed contours were used to estimate frequency peaks and troughs per call. We obtained means and standard error of the number of peaks per call, modulation rate (number of peaks divided by the duration of call), and the maximum slope of frequency modulation per call (largest negative slope between peaks and neighboring troughs). The effect size of range was calculated as Cohen’s *d* on the 3 frequency modulation measurements, as well as 15 standard acoustic measurements. Acoustic measurements with the largest effects of range were also compared among invasive calls sampled over time, to assess whether the patterns identified held over 15 years of sampling in the invasive range. Invasive range populations that grew over time could experience greater selection for more distinctive individual signatures, confounding direct comparisons between ranges.

### Individual identity content

Beecher’s statistic (HS) was employed to quantify the amount of individual identity content in calls of repeatedly sampled individuals per range [17]. We used 5 individuals per range, recorded at a single site-year in the native range (site 1145 in 2017), or a single city-year (Austin, TX in 2019) in the invasive range. These individuals showed similar patterns of dispersion in acoustic space (Supplementary Figure 2). HS was calculated by principal components analysis on Mel-frequency cepstral coefficients (MFCC) of all calls per individual, or the second harmonic frequency contours for 5 randomly sampled calls per bird (or all calls if 5 or less were recorded), with 5 points dropped on either end, and without spline interpolation. The number of possible unique individual signatures given the amount of individual identity content per range was estimated as 2^HS^ [9,17].

**Figure 1.**
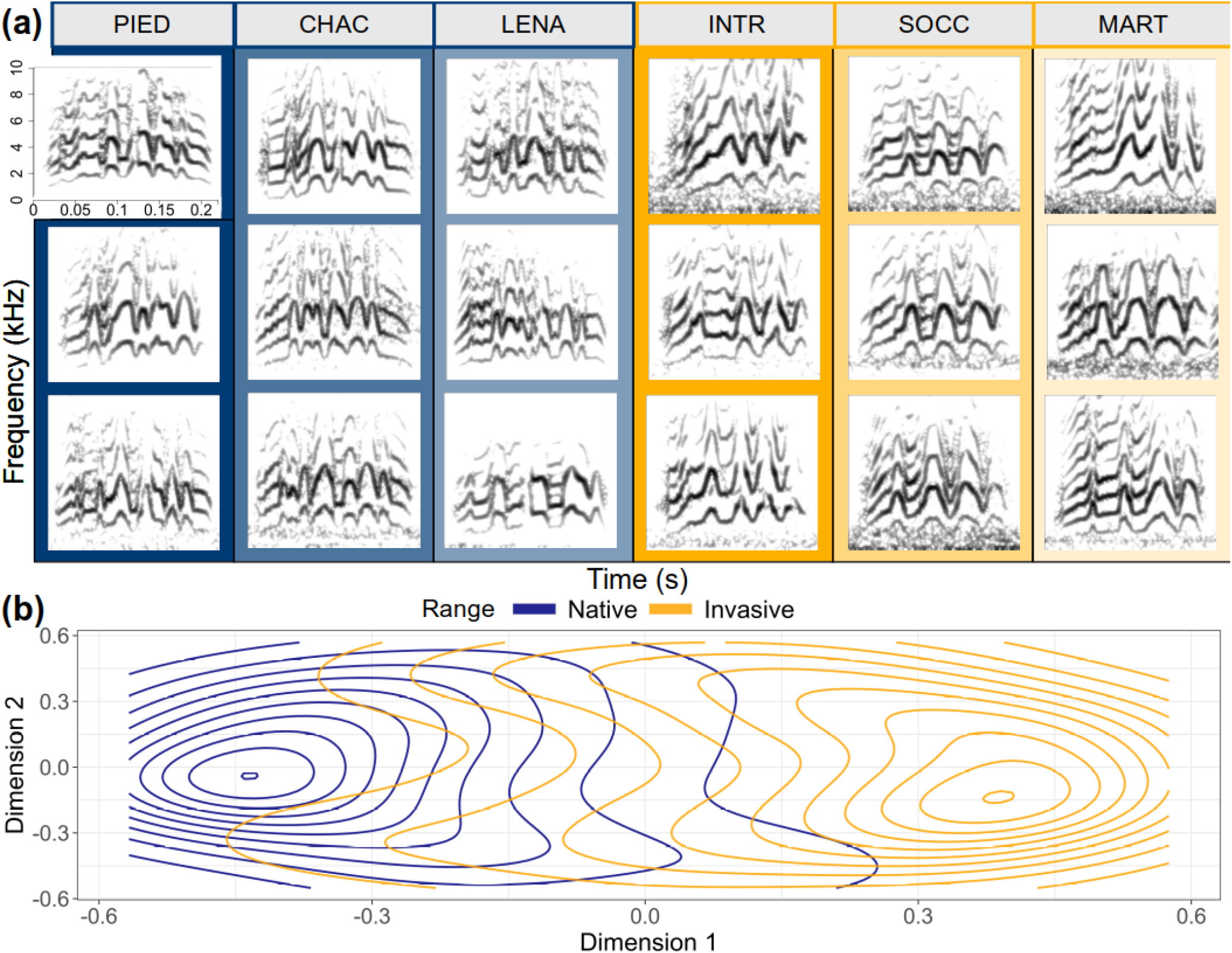
Differentiation in contact call structure between ranges. (a) Example lexicon with spectrograms of 3 randomly selected calls from 3 sites per range, each call represents a different unmarked individual. Calls were sampled over similar areas per range. Native and invasive range calls shown were recorded in 2017 and 2019, respectively. (b) Estimated kernel density contours in random forests acoustic space for the prediction dataset. Contours delineate bins of density values, with each bin representing 1/10th of the density per range. Dimension 1 coordinates were flipped to place native range contours on the left-hand side.

**Figure 2.**
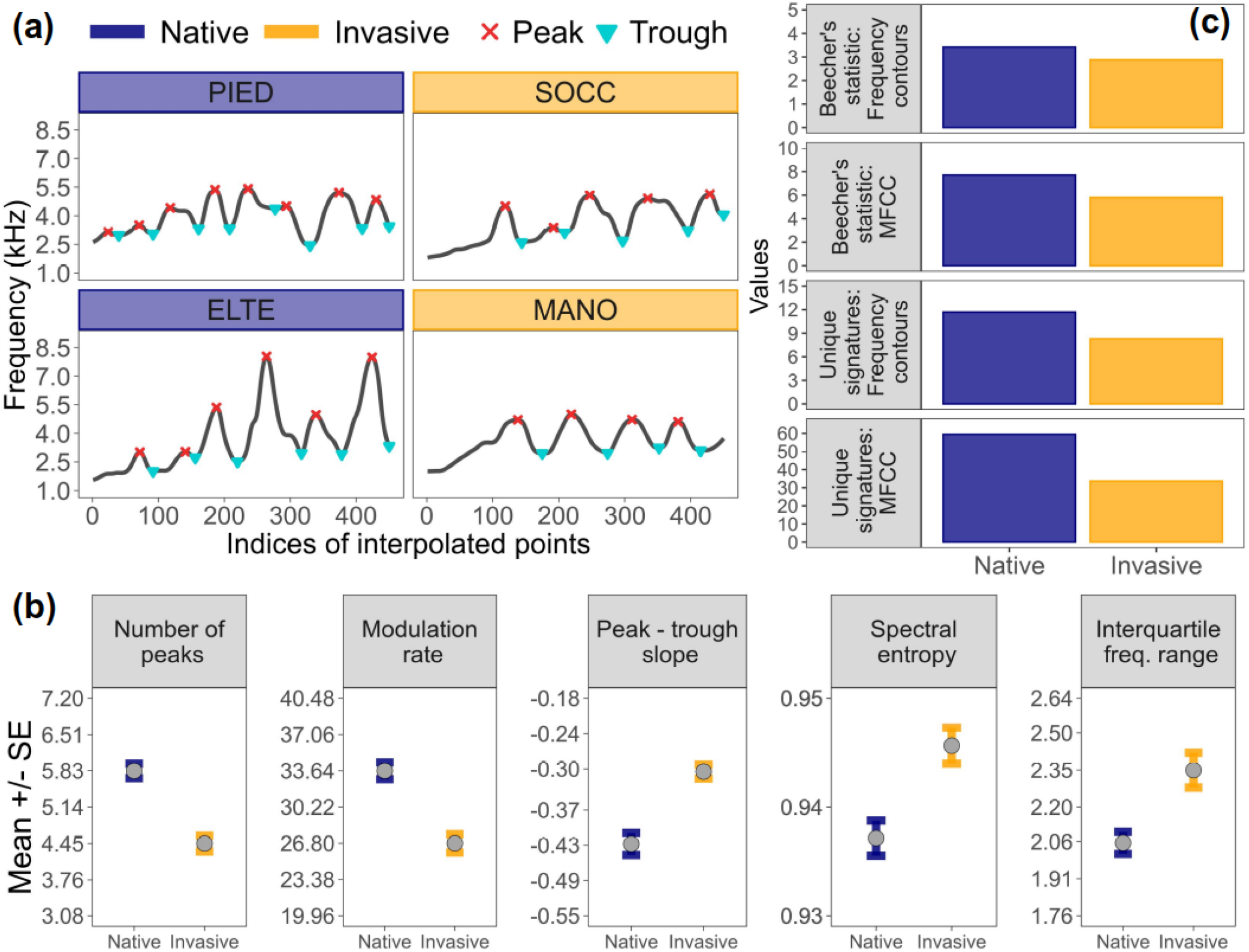
Simpler frequency modulation patterns in the invasive range. (a) Smoothed second harmonic frequency contours, marked with estimated peaks and troughs. (b) Mean and standard error of acoustic measurements with the largest effects of range, in decreasing order of absolute effect size magnitude (left to right). (c) Beecher’s statistic and the number of possible unique individual signatures, calculated with frequency contours as well as Mel-frequency cepstral coefficients.

## Results

### Lower nest density in invasive range

We observed (mean ± standard error) 36.35 ± 12.24 nests per site for the native range, and 5.94 ± 1.23 for the invasive range. The maximum number of nests observed at a given site was an order of magnitude greater in the native range (Supplementary Table 1), and nest estimates were significantly different between ranges (Mann-Whitney difference in location with 95% CI: 14 (7, 26), Z = 4.21, p < 0.0001).

### Simpler frequency modulation patterns in invasive range calls

Native and invasive range calls exhibited structural differences. Frequency modulation patterns were visibly different between ranges (Figure 1a), consistent with high classification accuracy by supervised random forests (Supplementary Table 5), and differentiation between ranges in random forests acoustic space (Figure 1b). Frequency modulation patterns contributed significantly to structural differences between ranges. Invasive range calls exhibited fewer frequency modulation peaks, lower modulation rates, and shallower maximum peak – trough slopes (Figure 2b). The effects of range on frequency modulation measurements were large and significant (Supplementary Table 6), and these trends were consistent over the 15 year sampling period in the invasive range (Supplementary Figure 1).

### Less individual identity content in invasive range calls

Beecher’s statistic was lower for invasive range calls. This reduced individual identity content yielded fewer distinctive individual signatures compared to native range calls, and trends were similar between MFCC and frequency contours (Figure 2c, Supplementary Table 7). MFCC includes frequency modulation patterns as well as other aspects of acoustic structure, including timbre and absolute frequency, that may arise from individual differences in vocal morphology.

## Discussion

Social group sizes and contact calls of native and invasive range monk parakeets were compared to ask whether the use of vocal learning for individual recognition could be sensitive to changes in the social environment following invasion. We found smaller social groups at invasive range sites, and frequency modulation patterns, which can be altered by learning and used for individual vocal recognition [4–6,8], were significantly simpler and contained less individual identity content in invasive range calls. Our results suggest that monk parakeets use vocal learning for individual recognition, and that this use of vocal learning for social recognition is sensitive to social changes associated with invasion.

Simpler frequency modulation patterns in the invasive range may be due to lower social densities and hence relaxed selection for individual vocal distinctiveness. Smaller invasive population sizes are a well-documented outcome of invasion [10]. Indeed, we observed fewer nests at invasive range sites, indicative of reduced local social densities compared to the native range. We do not know whether social dynamics are also altered following invasion, but this seems plausible given reduced population sizes as well as increased population isolation compared to the population contiguity observed in the native range (Smith-Vidaurre, pers. obs.).

Alternatively, the structural changes we identified could be due to a withdrawal of learning founder effect [18–20], if invasive populations were established by juvenile and/or captive birds that lacked adult tutors and thus developed atypical calls. We consider this alternative less likely, because changes in acoustic structure were concentrated on aspects of frequency modulation rather than distributed across all acoustic measurements. Furthermore, these changes were seen across multiple, presumably independent, invasions in 5 different states. Another possibility is that structural change could be due to genetic bottlenecks, another common outcome of biological invasions [21,22]. If structural variation in contact calls had a strong genetic component, acoustic variation should have decreased in concert with the reduced neutral genetic variation previously reported in the U.S. [23], yielding high overlap among individuals in acoustic space over short geographic distances. Instead, we identified similarly high levels of acoustic variation among individuals in the invasive range as previously found in the native range, indicating that although invasive range calls contain less individual identity information, individuals in both ranges are using learning to diverge in acoustic space (Smith-Vidaurre et al., unpublished data).

Reduced individual identity content in invasive range calls is consistent with relaxed selection for individual recognition in smaller populations. Beecher’s statistic (HS) calculated with Mel-frequency cepstral coefficients (MFCC) estimated 59 unique individual signatures for the native range versus 33 for the invasive range, while HS from frequency contours estimated 11 versus 8 unique signatures. Future work should manipulate social group size as well as social dynamics to ask whether vocal learning facilitates altering individual signatures to match changes in social group complexity over short timescales. Monk parakeets exhibit high fission-fusion fluidity [24], but how social dynamics influence individual recognition remains an open question.

Simpler individual signatures may be favored due to cognitive costs, or may arise from developmental constraints. In larger social groups, receivers incur the cognitive costs of discriminating among more individuals while simultaneously processing more complex individual signatures against a noisier background. Signalers should also experience costs of learning to encode more distinctive individual signatures through fine-scale structural variation. Monk parakeets both produce and recognize contact calls, therefore all individuals should experience costs of both receiver perception and vocal production [3]. As the perception and production of complex individual signatures impose greater cognitive burdens, simpler individual signatures should be present in smaller groups in which accurate recognition is possible without such complexity. Simpler or less informative signaler traits should also be favored when the costs of errors in individual recognition are lower [25]. Both factors could be working in the smaller populations found in the invasive range to yield simpler signatures. Finally, simpler individual signatures may reflect developmental constraints in receiver perception. Although parrots are considered open-ended vocal learners [26], we do not know whether auditory perception remains sensitive throughout adulthood [27]. In monk parakeets, perception of individual signatures may be constrained by local social densities experienced during sensitive developmental periods. The findings we present here on complexity of individual signatures produced by an invasive parrot add to a foundation for future work on how vocal learning is employed for individual recognition in dynamic social groups.

## Acknowledgments

We thank Clara Hansen for help trapping, marking and recording parakeets in Uruguay as well as Tania Molina for help with Uruguay fieldwork. We thank numerous additional people for their support throughout native range fieldwork, as acknowledged in Smith-Vidaurre et al. (2020). We are grateful to Zoë Amerigian and Alexandra Bicki for their help during 2019 fieldwork, and Susannah Buhrman-Deever for providing data. This manuscript benefited from discussion with Dr. Alejandro Salinas-Melgoza, Dr. Elizabeth Hobson, and Dominique Hellmich.

