## Supplementary Methods for "Individual vocal signatures show reduced complexity following invasion"

**Supplementary Materials:**

R code and RMarkdown output are available on GitHub (http://gsvidaurre/simpler-signatures-post-invasion). Pre-processed data will be deposited in Dryad. Included below are supplementary methods, tables, and figures.

**Supplementary Methods:**

*1. Contact call recording and pre-processing*

*1.1 Recording calls and obtaining nest estimates*

Contact calls were recorded similarly across ranges and years. Calls were generally obtained from unmarked parakeets flying in or out of clusters of nests, as well as perched individuals, as in [1,2]. With the exception of a subset of individuals, we obtained a single call per unmarked bird. As birds were unmarked, some calls may represent potential repeated sampling of the same individuals. Recordings were made using recording rigs, sampling rates and bit depths detailed in the main manuscript. Recordings were made onto a single channel. The 2004 calls provided as cuts of original recordings were previously high-pass filtered at 600Hz to remove low frequency noise in the background [1].

Numbers of nests were estimated at some native range recording sites in 2017, and some invasive range sites in 2011 and 2019 (Supplementary Table 1) by counting the number of nests visible at each site. Numbers of nests reported here should be considered estimates because other nests in the vicinity may have been missed, it was not always possible to evaluate if nests were active, and we could not always count the number of chambers, nor the number of individuals residing in each chamber. Overall, we observed greater numbers of parakeets and population continuity in the native range compared to invasive range sites in the U.S. (Smith-Vidaurre, pers. obs.). Although such native range numbers and continuity was not fully captured by nest estimates reported here, we used estimated numbers of nests as a rough proxy of local social density per range.

The effect size of range on nest estimates was calculated as Cohen's d with the effsize package version 0.8.0 with 95% CI : -0.75 (-0.06, -1.44). We asked whether nest estimates were significantly different between ranges with a Mann-Whitney-Wilcoxon test, as data were not normally distributed. To meet the assumption of independent samples, 4 invasive range site-years were dropped that represented sampling of the same sites over two years (sites AIRP, ELEM, INTR, MART in 2011 were dropped). This yielded 33 nest estimates for unique sites across ranges, with similar means and standard error for the invasive range as for the full dataset (reduced dataset: 5.31 ± 1.09, full dataset: 5.94 ± 1.23). The Mann-Whitney-Wilcoxon test was carried out with the package coin version 1.3-1 as a two-sided test. The distributions of nest estimates were not equal between ranges. The difference in location between ranges and 95% CI was 14 (7, 26), with $Z = 4.21$ and $p = 0.0000029$. The positive sign of this shift was consistent with greater nest estimates in the native range.

### *1.2 Call selection in Raven and pre-processing calls in R*

Contact calls were manually selected in Raven version 1.5 [3] from 2017 native range recordings in previous work [4]. Calls were selected from 2011, 2018, and 2019 invasive range recordings with Raven 1.4 [3]. Previously published 2004 contact calls were provided as cuts of original recordings [1]. Unless specified otherwise, call pre-processing was performed in R version 3.4.4 [5] with the warbleR package version 1.1.18 [2]. Invasive range calls, including 2004 calls, were taken through a similar pre-processing workflow as in [4]. We made catalogs of invasive range calls and visually checked call quality. Calls were assigned a score of low, medium or high visual quality. We also checked for visible patterns of amplitude saturation, overlapping signals in the background, and visible truncation of calls (2004 cuts), and added this metadata to a spreadsheet for manually detected calls. We used this metadata to retain high quality calls. Calls with low quality scores, visible amplitude saturation, overlapping signals, or signal to noise ratio less than 7 were dropped, as in [4].

Temporal coordinates of calls were tailored by the same observer (GSV, who tailored native range temporal coordinates in previous work) to return consistent start and end times across the native and invasive range datasets. Spectrograms were generated for individual calls to visually validate call quality and consistency of temporal coordinates, using the following settings: Hanning window, window length of 398, window overlap of 90. Unless otherwise specified, we used the same settings for all measurements below relying on Fourier transformations (e.g. spectrographic cross-correlation), in addition to a bandpass filter of 0.5 to 9kHz. Native and invasive range selection tables were combined, and filtered to retain sites with 5 calls or more remaining after pre-processing (Supplementary Tables 2, 3), and repeatedly sampled individuals with 4 or more calls (Supplementary Table 4).

We dropped duplicate recording sessions when a site was re-recorded on different days. However, some sites in the current dataset were represented by calls recorded on different days. This was due to merging sites that represented very fine-scale geographic sampling, which had been used for previous comparisons of geographic variation in the native range [4] (Supplementary Table 2). Also, for an independent analysis of hierarchical mapping patterns, we included 1 call per repeatedly sampled individual at the site scale, which led to more than one recording date for some sites with known repeatedly sampled individuals. The full dataset contained 1596 calls across social scales (individual scale = repeatedly sampled individuals, site scale = 1 call per “unique” individual) and ranges. However, for supervised machine learning analyses below, we dropped calls of repeatedly sampled individuals included at the site scale to avoid including duplicated calls, yielding a total of 1561 calls. See the script "SimplerSignatures_AdditionalMaterials_01_SummaryStatistics.Rmd" and the RMarkdown output provided on GitHub for more information.

*2. Analyses of acoustic structure*

*2.1 Supervised machine learning classification*

*2.1.1 Obtaining predictors for machine learning*

We measured a large set of acoustic measurements, including a standard set of 27 acoustic measurements and Mel-frequency cepstral coefficients (MFCC). Acoustic similarity of calls was measured using spectrographic cross-correlation (SPCC), dynamic time warping (DTW) on spectral entropy and dominant frequency time series estimated at 100 timepoints per call, and multivariate DTW (multiDTW) on spectral entropy and dominant frequency time series. These acoustic and similarity measurements were calculated with warbleR version 1.1.18 in R version 3.4.4. Acoustic measurements were converted to features for supervised machine learning using principal components analysis (PCA), and similarity measurements were converted to features via multidimensional scaling (MDS). Converting raw measurements to features yielded new predictors that represented variation across calls while reducing collinearity present among the original raw measurements.

We filtered out calls from the site scale that represented repeatedly sampled individuals included for a separate analysis of hierarchical mapping, yielding 1561 calls for supervised machine learning analyses (see section 1.2). We combined features extracted with MDS and PCA (see above) with 27 standard acoustic parameters, yielding 217 predictors. This set of predictors was filtered for high collinearity using Pearson's correlation (predictors with Pearson's *r* less than or equal to 0.75 were retained). After dropping highly collinear predictors, we obtained a final set of 203 predictors for machine learning, which included 15 acoustic measurements (see below), and 188 features derived by MDS and PCA. The 15 acoustic measurements were: start and end dominant frequency, minimum and maximum dominant frequency, dominant frequency range and slope, modulation index (based on dominant frequency), peak frequency, mean peak frequency, frequency interquartile range, third frequency quartile, kurtosis, spectral entropy, duration, and first temporal quartile. These acoustic measurements were used as predictors so as to directly evaluate their importance for classification of calls back to ranges, as it is easier to attribute structural differentiation to original measurements (such as call duration) rather than features representing less interpretable combinations of original measurements (e.g. principal components).

*2.1.2 Splitting calls for machine learning*

We split the dataset of 1561 calls into training, validation, and prediction datasets in R version 3.6.3. All subsequent analyses were performed with this version of R. Calls per site were randomly split depending on whether or not a site was used for spatial or temporal comparisons of acoustic structure. For native range sites and each invasive range site that did not represent temporal sampling, we randomly sampled ½ of total calls for training. Among the remaining calls per site, we randomly sampled 1/3 for validation, and set aside the rest (2/3) for prediction. For invasive range sites that did represent temporal sampling (e.g. the same site sampled in different years, or sites representing a city sampled over years, only Austin, TX and New Orleans, LA sites), we randomly sampled 20 calls for prediction. If one of these sites had 20 calls or less, we took all calls for prediction. For temporally sampled sites with more than 20 calls, we randomly chose ½ of the remaining calls for training, and set aside the other half for validation.

This overall sampling scheme yielded 676 calls for training, 337 calls for validation and 548 calls for prediction, while sampling as evenly as possible from different spatial regions and years in the invasive range dataset. Training, validation, and prediction datasets contained 43%, 22%, and 35%, respectively, of all calls used for supervised machine learning. The prediction dataset contained invasive range calls from all areas sampled in the U.S. for our direct comparison between ranges, and also contained invasive range calls sampled over time in Austin and New Orleans to assess the possibility of structural change in invasive range calls over time.

*2.1.3 Model training, validation, and prediction*

Supervised stochastic gradient boosting and random forests models were built to classify calls back to either the native or invasive range. Models were trained and tuned with the 203 predictors described above over 5 iterations of repeated 5-fold cross-validation using caret version 6.0-86, gbm version 2.1.5, and ranger version 0.12.1. The total number of trees, interaction depth (maximum depth of each tree, or the highest level of interactions permitted among predictors) and shrinkage parameter (learning rate of the model) were tuned for the gradient boosting model. The mtry parameter (the number of predictors randomly selected at each split) was tuned for the random forests model. After evaluating training performance, we visualized variable importance per model. Although the random forests model had slightly lower training classification accuracy, it exhibited more original acoustic measurements among the top 30 most important variables for classification back to ranges. As we wanted to use these acoustic measurements to more closely evaluate structural differences between ranges, we selected the random forests (RF) model for validation and prediction. This model yielded high validation accuracy, so we proceeded with prediction, and found that the model demonstrated high prediction accuracy back to ranges (Supplementary Table 5).

*2.1.4 Finer-scale assessment of structural change*

High classification accuracy during model training, validation, and prediction indicated high structural differentiation between ranges. These structural differences were visualized by reducing the RF proximity matrix to two dimensions with MDS. Density in acoustic space per range was obtained by applying a two-dimensional Gaussian kernel density estimator with bandwidth of 0.5 in each dimension to the MDS coordinates. Contours were drawn by splitting density values into 10 bins, such that each contour represented 1/10th of the density values per range (Figure 1b). Finer-scale spatial and temporal structural changes were evaluated by assessing classification accuracy of calls set aside for spatial and temporal comparisons in the RF prediction dataset, using misclassification back to the native range as an indicator of structural change (e.g. invasive range calls becoming more native range-like).

We expected that if invasive range populations grew in size over time, these populations should experience greater selection for more distinctive individual signatures, and therefore, invasive range calls could become more structurally similar to native range calls over time. If so, we expected to see higher misclassification of invasive range calls over time, or in different sampling areas that may have exhibited larger population sizes but were not sampled over time. However, we found no clear changes in classification accuracy over regions or years in the invasive range, which indicated that structural differences identified between ranges largely held regardless of the year and region in which invasive populations were sampled (see code provided). We also validated misclassification of invasive range calls and found that misclassification was not due to low signal to noise ratio (e.g. misclassified calls were not lower quality calls). Finally, structural changes in calls between ranges were assessed at a finer structural scale by assessing partial dependency of RF classification accuracy on the 15 standard acoustic measurements used among predictors. Partial dependency plots showed little change in classification accuracy back to the invasive range, indicating that structural differences between ranges did not entirely map onto these 15 standard acoustic measurements. See the script "SimplerSignatures_AdditionalMaterials_02_AcousticStructure_SupervisedML.Rmd" and the RMarkdown output provided on GitHub for more information.

*2.2 Obtaining second harmonic frequency contours*

*2.2.1 Randomly selecting calls for three comparisons (between ranges, over time, among individuals)*

The full dataset of 1596 calls was subsampled for frequency tracing, as we relied on manual tracing and this would have been prohibitively time-consuming to perform for the entire dataset. We randomly selected a subset of calls from the site scale dataset (e.g. not the dataset of known repeatedly sampled individuals) for a spatial comparison between the native and invasive ranges, as well as calls for a temporal comparison within the invasive range. We used temporal comparisons to account for the possibility of temporal change in acoustic structure, which could confound direct comparisons between ranges. 10 sites were randomly selected per range, and 4 calls randomly chosen per site. Overall, 80 calls were selected to evaluate frequency modulation patterns between ranges. These calls represented all sampling regions in the native range relatively evenly, although Texas was more heavily represented in the invasive range calls, as the full dataset contained more calls from this area.

For temporal comparisons of frequency modulation, we chose 15 site-years from Austin and New Orleans that represented sampling over time. Austin sites were each sampled in two years (10 site-years total, sampled in either 2011 and 2019, 2004 and 2019, or 2004 and 2011), while the same New Orleans sites were not sampled over different years, but together represented temporal sampling at the city scale (3 sites sampled in 2004, 2 sites sampled in 2011). We randomly selected 5 calls per each site-year, yielding a total of 75 invasive range calls for temporal comparisons. 25 calls were selected for 2004 (Austin and New Orleans), 30 calls represented 2011 (Austin and New Orleans), and 20 calls were sampled for 2019 (Austin only).

We also randomly sampled 5 calls per repeatedly sampled individual per range, or took all calls for repeatedly sampled individuals with 5 calls or less, yielding a total of 84 calls used for analyses of individual identity content (section 3.1.2). Our overall sampling scheme for frequency tracing yielded 239 calls total, but 6 calls were randomly sampled from 3 site-years for both the spatial and temporal comparisons (1 call from BALL-2004 in New Orleans, 1 call from INTR-2011 in Austin, and 4 calls from VALL-2004 in Austin), so we performed frequency tracing for 233 calls total.

*2.2.2 Tracing second harmonic frequency contours*

Frequency contours were obtained by estimating fundamental frequency as a time series at 100 timepoints per call, and these contours were used to manually trace the second harmonic per call with warbleR version 1.1.24 [2]. Unless otherwise specified, we used this version of warbleR for all subsequent analyses. We chose to trace the second harmonic because the fundamental frequency was not always clearly visible across calls. The subset of calls selected above for frequency tracing was randomly split in half to spread the manual tracing workload across two observers (GSV, VP). Tailored contours per observer were then combined, and a final round of tailoring was performed by one observer (GSV). Finally, spectrograms of calls with frequency contours were generated and inspected as a final check of tracing accuracy, and frequency contours were saved in extended selection table format.

*2.3 Frequency modulation analyses*

*2.3.1 Estimating peaks and troughs of frequency contours*

To measure frequency modulation patterns, we dropped 5 points from the start and end of each contour to account for small gaps preceding or following calls, and some end points that fell underneath components of the graphical user interface used for tailoring. We then randomly selected 5 calls per range from the subset of calls with frequency contours and generated image files of the frequency contours. One observer (GSV) manually counted large, visible frequency peaks and troughs per call. This step was performed in order to inform our approach for estimating peaks and troughs (inverted peaks). Once we obtained the number of visible peaks and troughs per call, we applied a general peak locating function to frequency contours of the randomly sampled set of 10 calls above, using pracma version 2.2.9. This initial peak search was used to fine-tune a more customized peak and trough estimation routine across the 233 calls with frequency contours.

From the preliminary peak search above, we obtained the maximum peak height identified in the subset of 10 calls, and used this to implement a threshold on minimum peak height in the customized function below. We also implemented smoothed spline interpolation of frequency contours, using the built-in R package stats version 3.6.3. Spline interpolation was performed with an exact cubic spline over 5 times the length of each frequency contour (e.g. 450 points), with means obtained for tied values. Cubic smoothing splines were applied to the interpolated points, and we optimized degrees of freedom, a parameter that controlled the degree of smoothing. Spline interpolation and smoothing helped flatten small peaks introduced by manual tailoring. pracma was used as above to estimate frequency peaks of the smoothed spline-interpolated points. Limitations were imposed on the peaks identified by pracma: peaks could not be within 2 points of the end of the smoothed frequency contour, peaks had to exhibit heights greater than a minimum height threshold (obtained above) compared to preceding frequency points, and peaks had to be a minimum distance apart (to filter out multiple peaks identified when a single tall peak presented as a plateau). We estimated troughs by searching for peaks across the inverted smoothed contours with pracma. Once troughs were obtained, troughs were assigned to closest preceding peaks, and we removed troughs that were not assigned to peaks. This routine returned peaks and troughs per call, as well as the slope per peak – trough pair (change in frequency/change in indices of smoothed contours), and image files for visual inspection of results.

We applied this customized function to the 233 calls with frequency contours, and visually inspected the peaks and troughs estimated per call to settle on final parameters for the function. Overall, the customized peak – trough estimation routine performed well when estimating large frequency peaks, and identifying troughs following each large peak. In a few cases, medium or small frequency peaks close to large peaks were not identified, and in other cases, gradual increases in frequency were labeled as peaks (and sometimes were not assigned troughs). Missing peaks per call could lead to underestimation of frequency modulation measurements. However, we felt this would not bias our results because peaks were missed for only a few calls in the dataset, and when this did occur, only a single peak was missed per call. In addition, the peaks missed were of small/medium height, and not representative of large changes in frequency modulation. On the other hand, visual inspection indicated that overestimation of frequency modulation was more of a problem (very small peaks or gradual rises in frequency identified as peaks). We addressed this concern by 1) removing peaks per call that were not matched to troughs, and 2) binning peak-trough slopes into 50 classes and removing peaks in the last two bins, which represented very small or positive peak-trough slopes. After dropping 170 peaks in these two bins, we proceeded with frequency modulation measurements across the 233 calls.

*2.3.2 Frequency modulation measurements*

Frequency modulation patterns were assessed by obtaining three frequency modulation measurements: the total number of peaks, the modulation rate (number of peaks/call duration), and the maximum peak – trough slope (largest negative slope between a given peak and neighboring trough) per call. We compared means and standard errors for each frequency modulation measurement per range, as well as for the 15 standard acoustic parameters filtered for high collinearity that were previously used for machine learning (section 2.1.1), with the set of 80 subsampled calls as described above. The effect size of range was calculated as Cohen's *d* on the 18 acoustic measurements, with pooled standard deviation and 95% CIs, using effsize version 0.8.0. We used Cohen's rule of thumb to identify large effect sizes, such that absolute effect sizes greater than or equal to 0.8 were considered large [6], and treated effect sizes with 95% CIs that did not cross zero as statistically significant (Supplementary Table 6).

We accounted for the possibility of temporal change in acoustic structure for the invasive range by evaluating means and standard errors of the 5 acoustic measurements with the largest effect sizes in the comparison above between ranges. Here we used the dataset of 75 calls selected for temporal comparisons. There was little change over time in these 5 acoustic measurements, indicating that the structural differentiation we identified between ranges was consistent over sampling intervals in the U.S. that spanned 15 years (Supplementary Figure 1). See the script "SimplerSignatures_AdditionalMaterials_03_AcousticStructure_FrequencyModulation.Rmd" and the RMarkdown output provided on GitHub for more information.

*3. Assessing individual identity content*

*3.1 Validation analysis of individuals used to calculate Beecher's statistic*

We used Beecher's statistic to calculate the amount of individual identity content in calls of repeatedly sampled individuals per range [7]. Here, we felt it was important to use equal numbers of individuals that represented similar patterns of variation in acoustic space per range. Previous work indicated that individuals at the same nesting site, as well as nesting sites separated by short geographic distances, are over-dispersed in acoustic space, but individuals begin to overlap in acoustic space over increasing geographic distances [4]. In our individual scale dataset, the 3 native range sites at which we repeatedly sampled individuals were separated by greater distances (minimum distance of 11.12km apart) than the 3 sites sampled for the invasive range (3.44 – 7.45km apart), which we felt could influence Beecher's statistic if native range individuals separated by greater geographic distances overlapped more in acoustic space. Therefore, we identified 5 repeatedly sampled individuals that represented restricted geographic areas per range, recorded at either a single site-year in the native range (site 1145 in 2017), or recorded at 3 sites in single year (city of Austin in 2019) in the invasive range. As it was not possible to assess 5 repeatedly sampled individuals at a single site in the invasive range, we performed a validation analysis to ask whether these individuals indeed represented similar patterns of call variation per range.

A bootstrapping analysis was designed to evaluate patterns of variation in second harmonic frequency contours represented by three sets of individuals: 5 native range individuals randomly sampled from 3 sites, the 5 native range individuals recorded at a single site (1145), and the 5 invasive range individuals recorded in Austin 2019. DTW was performed on second harmonic frequency contours (no spline interpolation or smoothing, 5 points were dropped from the start and end of each contour) to obtain pairwise acoustic distances. Per bootstrapping iteration, we randomly sampled 4 calls per individual (or took all calls if there were only 4 total). For the native range comparison with 3 sites, we randomly sampled 5 of the 8 total individuals recorded over 3 sites. Then per individual, we obtained the difference in mean DTW distance within each individual compared to other individuals for the given range and comparison. This process was repeated over 1000 iterations. The mean difference in DTW distance and 95% CIs were calculated per range and comparison. Mean DTW differences were similar between the 5 native range individuals from a single site and the 5 invasive range individuals at 3 sites, but were lower for the 5 individuals randomly sampled from 3 native range sites (Supplementary Figure 2). Therefore, the individuals from the 3 native range sites (representing greater geographic spread than the invasive range individuals) were more likely to overlap in acoustic space. The native range individuals from a single site and the invasive range individuals from 3 sites did indeed represent similar patterns of acoustic variation, so we proceeded with these individuals for Beecher's statistic calculations.

*3.2 Calculation of Beecher's statistic*

Beecher's statistic (HS) was calculated through the IDmeasurer package version 1.0.0 [7] using two acoustic measurements: MFCC calculated from all calls per individual, and second harmonic frequency contours for 5 randomly sampled calls per bird (or all calls if 5 or less were recorded). As in frequency modulation analyses above, 5 points were dropped on either end of each frequency contour, but we did not perform spline interpolation or smoothing. HS was reported using the sum of principal components significantly related to individual identity (e.g. significantly different among individuals) (Supplementary Table 7). We estimated the number of potential unique individual signatures per range and measurement as $2^{HS}$ [7] (Supplementary Table 7). See the script "SimplerSignatures_AdditionalMaterials_04_IdentityContent.Rmd" and the RMarkdown output provided on GitHub for more information.

Supplementary Table 1: Nest estimates in the native and invasive ranges

| Range | Year | Department or City, State | Site Code | Estimated Nests |
|---|---|---|---|---|
| Native | 2017 | Maldonado | PLVE | 10 |
| Native | 2017 | Colonia | RIAC | 109 |
| Native | 2017 | San José | ECIL | 247 |
| Native | 2017 | Colonia | INES-01 | 10 |
| Native | 2017 | Colonia | SEMI | 29 |
| Native | 2017 | Colonia | INES-03 | 50 |
| Native | 2017 | Colonia | INES-07 | 15 |
| Native | 2017 | Colonia | INES-06 | 20 |
| Native | 2017 | Colonia | INES-08 | 25 |
| Native | 2017 | Colonia | INES-05 | 6 |
| Native | 2017 | Colonia | 1145 | 8 |
| Native | 2017 | Colonia | ROSA | 41 |
| Native | 2017 | Colonia | CHAC | 19 |
| Native | 2017 | Canelones | INBR | 20 |
| Native | 2017 | Montevideo | BCAR | 33 |
| Native | 2017 | Maldonado | HIPE | 15 |
| Native | 2017 | Maldonado | QUEB | 10 |
| Native | 2017 | Maldonado | CISN | 9 |
| Native | 2017 | Colonia | PIED | 38 |
| Native | 2017 | Rocha | VALI | 13 |
| Invasive | 2018 | Gilbert, AZ | GILB | 3 |
| Invasive | 2019 | Austin, TX | INTR | 13 |
| Invasive | 2019 | Austin, TX | ELEM | 1 |
| Invasive | 2019 | Austin, TX | AIRP | 5 |
| Invasive | 2019 | Austin, TX | SOCC | 12 |
| Invasive | 2019 | Austin, TX | MANO | 4 |
| Invasive | 2019 | Austin, TX | MART | 8 |
| Invasive | 2011 | Austin, TX | MART | 6 |
| Invasive | 2011 | Austin, TX | VALL | 2 |
| Invasive | 2011 | Austin, TX | ELEM | 4 |
| Invasive | 2011 | Austin, TX | SOFT | 6 |

| | | | | |
|---|---|---|---|---|
| Invasive | 2011 | Austin, TX | AIRP | 2 |
| Invasive | 2011 | Austin, TX | BART | 1 |
| Invasive | 2011 | Austin, TX | INTR | 20 |
| Invasive | 2011 | New Orleans, LA | ROBE | 8 |
| Invasive | 2011 | New Orleans, LA | LAKE | 2 |
| Invasive | 2011 | Dallas, TX | LAWT | 4 |

Supplementary Table 1 Footnote: Estimated numbers of nests for a subset of recording sites, ordered from most recent to later sampling years per range. Nest estimates were collected from Smith-Vidaurre, Perez, and Wright field notebooks.

Supplementary Table 2: Native range recording sites in Uruguay

| | Site Code | Site Name | Department | Latitude | Longitude | $N_{Calls}$ | Date |
|---|---|---|---|---|---|---|---|
| 1 | PIED | Piedra de los Indios | Colonia | -34.413 | -57.849 | 21 | 2017-10-25 |
| 2 | * CHAC | La Chacra de los Olivos | Colonia | -34.413 | -57.843 | 12 | 2017-08-21 |
| 3 | LENA | Las Leñas | Colonia | -34.411 | -57.838 | 19 | 2017-10-23 |
| 4 | PFER | Parque Ferrando | Colonia | -34.468,-34.465 | -57.831, -57.827 | 53 | 2017-06-19, 2017-06-21 |
| 5 | INES-08 | INIA La Estanzuela - 08 | Colonia | -34.345 | -57.733 | 27 | 2017-07-13 |
| 6 | * EMBR | Embarcadero de Riachuelo | Colonia | -34.444 | -57.728 | 23 | 2017-07-17, 2017-07-21 |
| 7 | INES-01 | INIA La Estanzuela - 01 | Colonia | -34.349 | -57.727 | 12 | 2017-07-03 |
| 8 | INES-07 | INIA La Estanzuela - 07 | Colonia | -34.346 | -57.710 | 9 | 2017-07-13 |
| 9 | INES-06 | INIA La Estanzuela - 06 | Colonia | -34.344 | -57.708 | 6 | 2017-07-13 |
| 10 | RIAC | Riachuelo | Colonia | -34.436, -34.437 | -57.706 | 25 | 2017-06-28 |
| 11 | INES-05 | INIA La Estanzuela - 05 | Colonia | -34.340 | -57.690 | 6 | 2017-07-15 |
| 12 | SEMI | Semillero | Colonia | -34.326 | -57.680 | 11 | 2017-07-25 |
| 13 | INES-03 | INIA La Estanzuela - 03 | Colonia | -34.336 | -57.668 | 15 | 2017-07-11 |
| 14 | INES-04 | INIA La Estanzuela - 04 | Colonia | -34.335 | -57.668 | 9 | 2017-07-11 |
| 15 | ARAP | Las Termas del | Salto | -30.946 | -57.520 | 12 | 2017- |

| | | | | | | | |
|---|---|---|---|---|---|---|---|
| | | Arapey | | | | | 05-07 |
| 16 | * 1145 | Ruta 1 km 145 | Colonia | -34.375, -34.376 | -57.502, -57.500 | 17 | 2017-07-24, 2017-07-26, 2017-07-28, 2017-07-29 |
| 17 | ROSA | Rosario | Colonia | -34.338 | -57.336 | 15 | 2017-07-27 |
| 18 | ECIL | Ecilda Paullier | San José | -34.360, -34.361 | -57.060 | 17 | 2017-07-28 |
| 19 | PAVO | Arroyo Pavón | San José | -34.442 | -56.967 | 25 | 2017-10-17 |
| 20 | ARAZ | Balneario de Arazati | San José | -34.535 | -56.812 | 15 | 2017-11-03 |
| 21 | KIYU | Balneario de Kiyú | San José | -34.607 | -56.715 | 8 | 2017-11-03 |
| 22 | BAGU | La Baguala | Montevideo | -34.848 | -56.384 | 20 | 2017-10-09 |
| 23 | INBR | INIA Las Brujas | Canelones | -34.668 | -56.330 | 19 | 2017-09-03 |
| 24 | PEIX | Camino Peixoto | Montevideo | -34.765 | -56.279 | 19 | 2017-10-06 |
| 25 | BCAR | Bodegas Carrau | Montevideo | -34.788 | -56.223 | 13 | 2017-10-20 |
| 26 | FAGR | Facultad de Agronomía | Montevideo | -34.838 | -56.219 | 7 | 2017-09-05 |
| 27 | CEME | Cementerio Central | Montevideo | -34.913 | -56.187 | 6 | 2017-10-18 |
| 28 | GOLF | Club de Golf | Montevideo | -34.923 | -56.164 | 22 | 2017-11-20 |
| 29 | PROO | Parque Roosevelt | Montevideo | -34.855 | -56.022 | 12 | 2017-09-14 |
| 30 | PLVE | Plaza Venus, Piriápolis | Maldonado | -34.870 | -55.264 | 11 | 2017-05-21 |
| 31 | QUEB | Quebrada del Castillo | Maldonado | -34.834 | -55.260 | 16 | 2017-09-13 |

| | | | | | | | |
|---|---|---|---|---|---|---|---|
| 32 | CISN | La Laguna de los Cisnes | Maldonado | -34.861 | -55.150 | 28 | 2017-09-13 |
| 33 | SAUC | La Laguna del Sauce | Maldonado | -34.857 | -55.041 | 6 | 2017-09-12 |
| 34 | HIPE | Centro de Entrenamiento Hípico Punta del Este | Maldonado | -34.825 | -55.010 | 5 | 2017-09-12 |
| 35 | ELTE | El Tesoro | Maldonado | -34.889 | -54.863 | 23 | 2017-09-13 |
| 36 | VALI | Barra de Valizas | Rocha | -34.334 | -53.803 | 23 | 2017-11-16 |
| 37 | OJOS | Ojos de Agua | Rocha | -33.804 | -53.506 | 23 | 2017-11-16 |

<u>Supplementary Table 2 Footnote:</u> Native range recording sites and dates in Uruguay. Numbers of calls recorded per site are reported (610 total). Asterisks denote the three sites at which we repeatedly sampled marked or unmarked individuals for the individual scale. Recording sessions per site were typically performed in a single day. However, when assessing invasive range sites in Austin recorded in different years to harmonize site codes for temporal analyses (in which sites recorded relatively close to each other in different years were assigned the same site code), we also merged 2 pairs of native range sites that been kept separate in our previous analyses (PFER-01, PFER-03 and RIAC-01, RIAC-02) to represent very fine-scale geographic sampling [4]. RIAC-01 (8 calls) and RIAC-02 (17 calls) were recorded on the same day, but PFER-01 (19 calls) and PFER-03 (34 calls) recording sessions were from different days. Moreover, for an independent analysis of hierarchical mapping patterns, when calls were merged into a single extended selection table across ranges, we added a single call per repeatedly sampled individual to the site-scale dataset per range, for consistency with previous work. This pre-processing led to calls recorded on different days for sites EMBR and 1145. The suffix of these calls is “_site_scale”, so these can be easily identified and/or removed as needed in future work. See section 1.2 for more details, and Supplementary Table 4 for repeatedly sampled individuals.

Supplementary Table 3: Invasive range recording sites in the U.S.

| | Site Code | Site Name | City, State | Latitude | Longitude | $N_{Calls}$ | Date |
|---|---|---|---|---|---|---|---|
| 1 | GILB | Gilbert Town Square | Gilbert, AZ | 33.331 | -111.791 | 16 | 2018-04-09 |
| 2 | LAWT | Lawther Substation | Dallas, TX | 32.820 | -96.730 | 9 | 2011-02-20 |
| 3 | COMM | Austin Community College | Austin, TX | 30.404 | -97.705 | 11 | 2004-03-30 |
| 4 | INTR | University of Texas (UT) – Austin Intramural fields | Austin, TX | 30.316 | -97.719 | 15 | 2011-02-15 |
| 5 | * INTR | UT – Austin Intramural Fields | Austin, TX | 30.317 | -97.727 | 82 | 2019-08-08 |
| 6 | MANO | Manor Rd. | Austin, TX | 30.299 | -97.728 | 5 | 2019-08-09 |
| 7 | AIRP | Airport Boulevard | Austin, TX | 30.285 | -97.705 | 9 | 2019-08-07 |
| 8 | SOFT | McCombs Softball Field | Austin, TX | 30.281 | -97.725 | 14 | 2011-02-15 |
| 9 | SOCC | Soccer Field, César Chavez | Austin, TX | 30.272 | -97.767 | 77 | 2004-03-30 |
| 10 | * SOCC | César Chavez Fields | Austin, TX | 30.270 | -97.761 | 93 | 2019-08-09 |
| 11 | VALL | Pleasant Valley Rd. | Austin, TX | 30.261 | -97.711 | 5 | 2004-03-30 |
| 12 | VALL | Pleasant Valley Rd. & 7th | Austin, TX | 30.261 | -97.711 | 10 | 2011-02-15 |
| 13 | ELEM | UT Elementary School | Austin, TX | 30.260 | -97.718 | 12 | 2011-02-15 |
| 14 | * ELEM | UT Elementary School | Austin, TX | 30.260 | -97.718 | 61 | 2019-08-06 |
| 15 | MART | Sam L. Martin Middle School | Austin, TX | 30.253 | -97.731 | 14 | 2011-02-15 |
| 16 | MART | Sam L. Martin Middle School | Austin, TX | 30.251 | -97.731 | 50 | 2019-08-10 |
| 17 | LAKE | Lakeview Dr. | New Orleans, LA | 30.029 | -90.077 | 6 | 2011-02-18 |
| 18 | FOLS | Folse Dr. & Harris St. | New Orleans, LA | 30.027 | -90.205 | 10 | 2004-03-30 |
| 19 | * ROBE | Robert E. Lee Rd. | New | 30.021 | -90.069 | 24 | 2011-02- |

| | | | | | | | |
|---|---|---|---|---|---|---|---|
| | | | Orleans, LA | | | | 18 |
| 20 | BALL | Ballfield at corner of W. Esplanade & Oaklawn | New Orleans, LA | 30.013 | -90.132 | 26 | 2004-03-30 |
| 21 | CANA | Canal Blvd. | New Orleans, LA | 29.981 | -90.110 | 13 | 2004-03-30 |
| 22 | BAPT | Baptist Hospital | Miami, FL | 25.6878 | -80.338 | 40 | 2004-03-30 |
| 23 | BUCK | Buckingham Ave. | Milford, CT | 41.217 | -73.038 | 60 | 2004-03-30 |
| 24 | MEAD | Meadowside Rd. | Milford, CT | 41.210 | -73.071 | 28 | 2004-03-30 |
| 25 | SHAK | Shakespeare Theatre | Stratford, CT | 41.184 | -73.126 | 50 | 2004-03-30 |
| 26 | AUDU | Milford Audubon | Milford, CT | 41.176 | -73.102 | 17 | 2004-03-30 |

Supplementary Table 3 Footnote: Invasive range recording sites and dates in the U.S. Numbers of calls recorded per site are reported (757 total). Sites recorded in 2004 were previously published [1]. Specific recording dates were not provided with the 2004 call dataset, so we assigned a single date to all 2004 sites within the dates reported by Buhrman-Deever et al. (2007). Geographic coordinates are also approximate for all 2004 sites, as we obtained these by entering site names in Google Maps. Site codes were harmonized over time for Austin as described above (Supplementary Table 1). Asterisks denote the three sites at which we repeatedly sampled unmarked individuals for the individual scale. See section 1.2 for more details, and Supplementary Table 4 for repeatedly sampled individuals.

Supplementary Table 4: Repeatedly sampled individuals per range

| | Individual ID | Site Code | Site Name | Depart-ment or City, State | Lati-tude | Longi-tude | $N_{Calls}$ | Date |
|---|---|---|---|---|---|---|---|---|
| 1 | NAT-AAT | 1145 | Ruta 1 km 145 | Colonia | -34.376 | -57.500 | 12 | 2017-07-29 |
| 2 | NAT-UM1 | 1145 | Ruta 1 km 145 | Colonia | -34.375 | -57.502 | 25 | 2017-07-28 |
| 3 | NAT-UM2 | 1145 | Ruta 1 km 145 | Colonia | -34.375 | -57.502 | 23 | 2017-07-24 |
| 4 | NAT-UM3 | 1145 | Ruta 1 km 145 | Colonia | -34.375 | -57.502 | 5 | 2017-07-24 |
| 5 | NAT-UM4 | 1145 | Ruta 1 km 145 | Colonia | -34.376 | -57.500 | 13 | 2017-07-26 |
| 6 | NAT-UM5 | CHAC | La Chacra de los Olivos | Colonia | -34.413 | -57.843 | 7 | 2017-08-21 |
| 7 | NAT-RAW | EMBR | Embarcadero de Riachuelo | Colonia | -34.444 | -57.728 | 4 | 2017-07-17 |
| 8 | NAT-ZW8 | EMBR | Embarcadero de Riachuelo | Colonia | -34.444 | -57.728 | 8 | 2017-07-21 |
| 9 | INV-UM6 | ASCA | Ascarate Park | El Paso, TX | 31.754 | -106.405 | 25 | 2019-03-10 |
| 10 | INV-UM10 | INTR | University of Texas (UT) – Austin Intramural fields | Austin, TX | 30.317 | -97.728 | 6 | 2019-08-08 |
| 11 | INV-UM7 | ELEM | UT Elementary School | Austin, TX | 30.260 | -97.718 | 28 | 2019-08-06 |
| 12 | INV-UM9 | ELEM | UT Elementary School | Austin, TX | 30.260 | -97.718 | 5 | 2019-08-06 |
| 13 | INV-UM16 | SOCC | César Chavez Fields | Austin, TX | 30.270 | -97.761 | 8 | 2019-08-09 |
| 14 | INV-UM17 | SOCC | César Chavez Fields | Austin, TX | 30.270 | -97.761 | 5 | 2019-08-09 |
| 15 | INV-UM1 | BART | Bartholomew Park | Austin, TX | 30.305 | -97.695 | 23 | 2011-02-15 |
| 16 | INV-UM5 | ROBE | Robert E. Lee Rd. | New Orleans, LA | 30.021 | -90.069 | 20 | 2011-02-18 |
| 17 | INV-UM19 | CAME | Robert E. Lee & Cameron Rd. | New Orleans, LA | 30.022 | -90.065 | 12 | 2004-03-30 |

Supplementary Table 4 Footnote: Number of calls, recording locations ,and dates for known repeatedly sampled individuals per range (229 total calls). Each individual was recorded on a single day. Native range individuals (prefix “NAT” in the Individual ID column) were recorded in Uruguay in 2017, while invasive individuals (prefix “INV” in the Individual ID column) were recorded in the U.S in 2019, 2011 or 2004. Two individuals were recorded at sites not included in the site-scale datasets due to insufficient sampling: sites ASCA and BART. Site CAME in 2004 was close to the site labeled ROBE recorded in 2011, but we did not harmonize site codes to be the same over time at the individual scale. The recording date for individual INV-UM19 at CAME 2004 is an approximate date from previously published work [1].

Supplementary Table 5: Supervised machine learning performance metrics

| Model | Training accuracy (%) and 95% CI | Final parameters | Validation accuracy (%) | Prediction accuracy (%) |
|---|---|---|---|---|
| Stochastic gradient boosting | 92.28 (91.33, 93.16) | n.trees = 1600, interaction.depth = 3, shrinkage = 0.1, nminobsinnode = 1 | - | - |
| Random forests | 91.09 (90.08, 92.03) | mtry = 2, splitrule = gini, min.node.size = 1, n.trees = 2000 | 91.99 | 87.59 |

Supplementary Table 5 Footnote: Supervised machine learning analyses of structural differences between ranges. Models were trained to classify calls back to the native or invasive range. The random forests model was selected for validation and prediction.

Supplementary Table 6: Effect sizes of range with 95% CI for 18 acoustic measurements

| | Measurement | Effect size | 95% CI |
|---|---|---|---|
| 1 | Number of peaks | 1.50 | **(2.00, 0.99)** |
| 2 | Modulation rate | 1.30 | **(1.79, 0.81)** |
| 3 | Peak – trough slope | -1.23 | **(-0.75, -1.72)** |
| 4 | Spectral entropy | -0.83 | **(-0.36, -1.30)** |
| 5 | Frequency interquartile range | -0.81 | **(-0.34, -1.28)** |
| 6 | Mean peak frequency | 0.57 | **(1.03, 0.11)** |
| 7 | Modulation index | 0.51 | **(0.96, 0.05)** |
| 8 | Dominant frequency range | -0.49 | **(-0.04, -0.95)** |
| 9 | Duration | 0.48 | **(0.94, 0.02)** |
| 10 | End dominant frequency | 0.44 | (0.90, -0.02) |
| 11 | Minimum dominant frequency | 0.42 | (0.87, -0.04) |
| 12 | Peak frequency | 0.40 | (0.85, -0.06) |
| 13 | First time quartile | 0.37 | (0.83, -0.08) |
| 14 | Third frequency quartile | -0.35 | (0.10, -0.81) |
| 15 | Kurtosis | -0.34 | (0.11, -0.79) |
| 16 | Maximum dominant frequency | -0.34 | (0.12, -0.79) |
| 17 | Start dominant frequency | 0.25 | (0.70, -0.21) |
| 18 | Dominant frequency slope | 0.15 | (0.61, -0.30) |

Supplementary Table 6 Footnote: Effect sizes of range on different acoustic measurements for 80 calls compared between ranges. Shown are 3 frequency modulation measurements and the 15 standard acoustic measurements used in supervised machine learning, in order of decreasing absolute effect size (top to bottom). Frequency measurements are in kHz and temporal measurements in seconds, although modulation rate is in peaks/s. Peak – trough slope represents change in kHz/change in indices of spline-interpolated points. 95% CIs that do not cross 0 (significant effect sizes) are in bold. Effect sizes greater than or equal to 0.8 were considered large [6]. Negative effect sizes indicate higher mean values for the invasive range, with the exception of peak – trough slope.

Supplementary Table 7: Beecher's statistic and possible unique individual signatures

| Acoustic measurements | Range | $N_{Calls}$ | HS | $N_{Sig}$ |
|---|---|---|---|---|
| 2nd harmonic | Native | 25 | 3.42 | 11.70 |
| | Invasive | 25 | 2.88 | 8.29 |
| MFCC | Native | 78 | 7.71 | 59.44 |
| | Invasive | 52 | 5.80 | 33.64 |

Supplementary Table 7 Footnote: Individual identity content in calls of repeatedly sampled individuals per range, using Beecher's information statistic (HS) with two measurements: Mel-frequency cepstral coefficients (MFCC) and second harmonic frequency contours. $N_{Sig}$ is the number of individual signatures predicted by HS. 5 individuals were used per calculation per range.

Supplementary Figure 1:

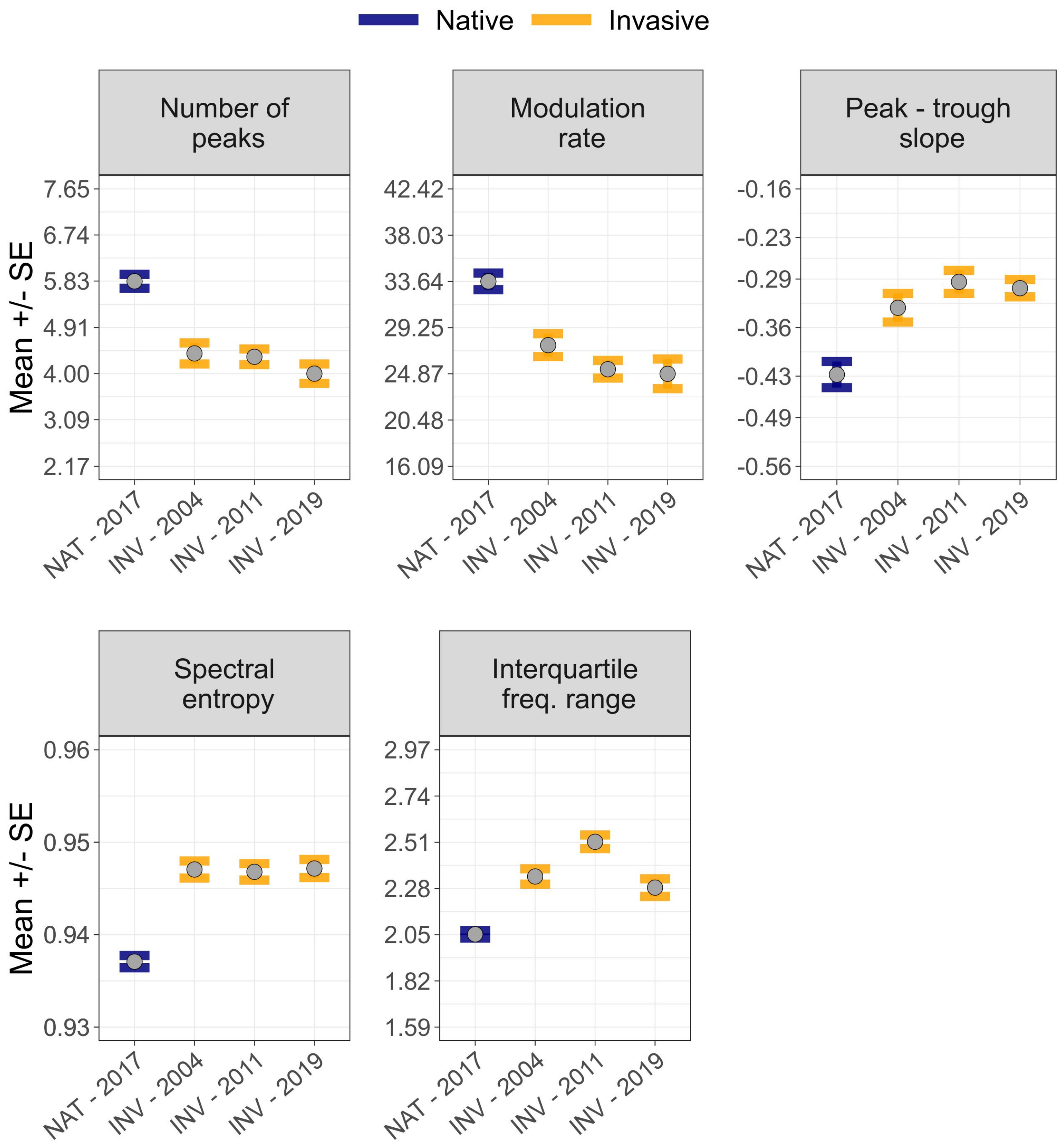


Supplementary Figure 1 Legend: Structural differences between ranges were stable over 15 years of sampling in the invasive range. Means and standard errors for the same acoustic parameters that displayed significant effects of range in Figure 2b. Invasive range-years represent 75 calls set aside for temporal comparison of frequency modulation measurements, and the 40 native range calls were used for the comparison between ranges.

Supplementary Figure 2:

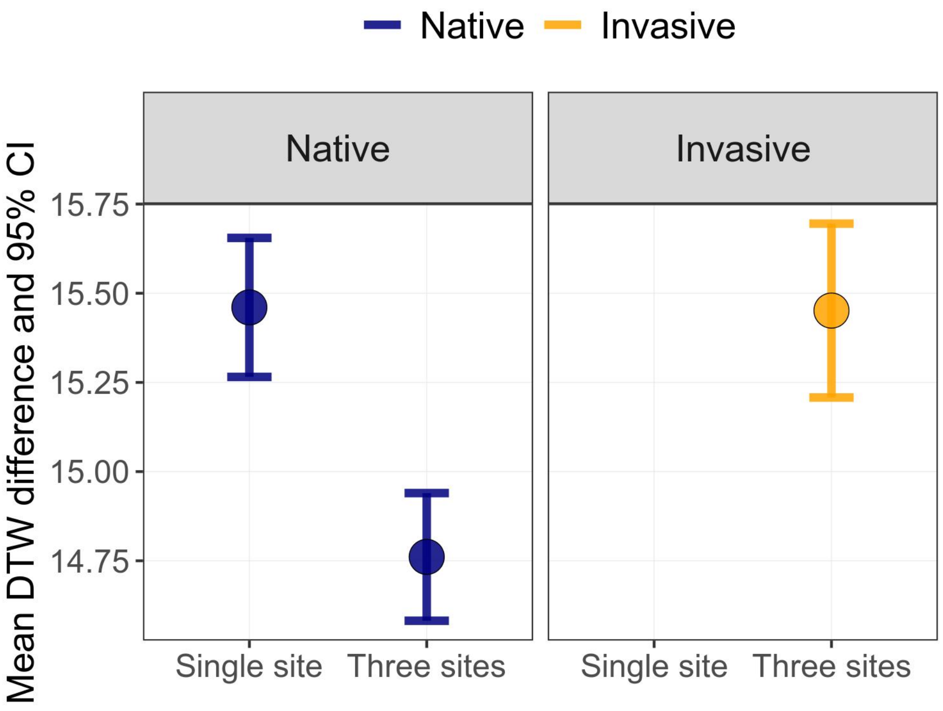


Supplementary Figure 2 Legend: Validation of repeatedly sampled individuals used for Beecher's statistic calculations. Shown are the mean differences in DTW distance of second harmonic frequency contours within an individual compared to among individuals at either a single site (native range only) or 3 sites (both ranges). The single site comparison is missing for the invasive range due to insufficient sampling of individuals. 5 individuals were used per comparison. 95% CIs were generated by bootstrapping with 1000 iterations. These results suggested that using 3 geographically proximate sites in the invasive range provided patterns of variation among individuals equivalent to using a single site in the native range, and supported using these individuals for direct comparisons of Beecher's statistic between ranges.
